# Differential effects of environmental and endogenous 24h rhythms within a deep-coverage spatiotemporal proteome

**DOI:** 10.1101/2021.03.30.437663

**Authors:** Holly Kay, Ellen Grünewald, Helen K. Feord, Sergio Gil, Sew Y. Peak-Chew, Alessandra Stangherlin, John S. O’Neill, Gerben van Ooijen

## Abstract

The cellular landscape of most eukaryotic cells changes dramatically over the course of a 24h day. Whilst the proteome responds directly to daily environmental cycles, it is also regulated by a cellular circadian clock that anticipates the differing demands of day and night. To quantify the relative contribution of diurnal versus circadian regulation, we mapped spatiotemporal proteome dynamics under 12h:12h light:dark cycles compared with constant light. Using *Ostreococcus tauri*, a prototypical eukaryotic cell, we achieved 85% coverage of the theoretical proteome which provided an unprecedented insight into the identity of proteins that drive and facilitate rhythmic cellular functions. Surprisingly, the overlap between diurnally- and circadian-regulated proteins was quite modest (11%). These proteins exhibited different phases of oscillation between the two conditions, consistent with an interaction between intrinsic and extrinsic regulatory factors. The relative amplitude of rhythmic protein abundance was much lower than would be expected from daily variations in transcript abundance. Transcript rhythmicity was poorly predictive of daily variation in abundance of the encoded protein. We observed coordination between the rhythmic regulation of organelle-encoded proteins with the nuclear-encoded proteins that are targeted to organelles. Rhythmic transmembrane proteins showed a remarkably different phase distribution compared with rhythmic soluble proteins, indicating the existence of a novel circadian regulatory process specific to the biogenesis and/or degradation of membrane proteins. Taken together, our observations argue that the daily spatiotemporal regulation of cellular proteome composition is not dictated solely by clock-regulated gene expression. Instead, it also involves extensive rhythmic post-transcriptional, translational, and post-translational regulation that is further modulated by environmental timing cues.

## Introduction

Endogenous circadian clocks drive organismal and cellular physiology over the 24 hours of the day/night cycle. Before the advent of ‘omics’ techniques, reverse genetic approaches identified “clock genes” in multiple organisms and facilitated the dissection of their auto-regulatory transcriptional-translational feedback loops (TTFLs) in the different taxonomic groups^1^. These feedback loops play a role in the regulation of circadian physiology, metabolism and many other aspects of cellular physiology^2^. Genetic and transcriptomic information are the major source of information upon which models of the cellular clock have been built.

However, abundant evidence also suggests that post-transcriptional and post-translational processes are essential to circadian regulation^3-5^. Indeed, in several different contexts, circadian rhythms have been observed in the complete absence of transcriptional feedback^6-11^. Since changes in protein activity underlie every biological process, these studies highlight the importance of studying eukaryotic circadian clocks at the proteome level when investigating the links between environmental signals, TTFLs and post-translational circadian regulation. However, functional circadian proteomics is currently in its infancy, compared with the detailed characterisation of circadian transcriptomes.

While temporally resolved proteomics datasets exist for a handful of model species (reviewed in^12^), methodological limitations have meant that the coverage of the theoretical maximum proteome is substantially lower than for transcriptomics. Lowly abundant proteins, cell cycle proteins, organelle-encoded proteins and transmembrane proteins tend to be underrepresented^13, 14^, as sample complexity exceeds the detection capacity of mass spectrometric analyses. This is reflected by a maximum of 45% total proteome coverage over circadian time series in the fungus *Neurospora*^14^, 30% in *Drosophila*^15^, 12% in *Arabidopsis*^16^, and 9% in mouse^17^. Furthermore, proteomics studies have exclusively been performed either under rhythmic environmental conditions (e.g. light:dark cycles) or under constant conditions. These experimental approaches are fundamentally different, as they either reflect the combined influences of environmental stimuli and circadian-regulated rhythms, or only the latter. Therefore, a direct comparison between different -omics studies is challenging since experimental details frequently vary^18^.

The aim of this study was to provide a detailed analysis of diurnal versus circadian proteomes in a single study, with unprecedented proteome coverage including organellar and integral membrane proteins. To reduce sample complexity, we employed the uniquely minimal cellular and genomic complexity of the model cell *Ostreococcus tauri*, a picoeukaryotic alga. This cell type is well established as a cellular model for circadian rhythms across eukaryotes^6, 9, 19^, and is highly amenable to culture under natural diurnal versus constant circadian conditions. Our results provide a rare insight into the complex relationship between environmental and circadian regulation of protein abundance across time, revealing a strikingly differential spatiotemporal proteome under these two conditions.

## Results

### A deep coverage diurnal and circadian proteome

To attain a deeper understanding of how cellular proteomes change over time, we established extraction procedures to enable increased coverage of transmembrane and organellar proteins in the minimal clock model system *Ostreococcus tauri*^20^ (Fig. 1A). We sampled under light-dark entrainment (LD) and constant circadian conditions (LL; Fig. 1B) and quantified the proteome by 11-plex Tandem Mass Tagging Mass Spectrometry (TMT-MS). This methodology allowed the detection of 86% of the 7700 nuclear-encoded proteins, of which 79% were detected at all time points (Fig. 1C and Supplemental File 1). ∼24 h rhythmicity was detected in 54% of proteins under LD (Fig. 1D), and 11% under LL (Fig. 1E). The observation that the *Ostreococcus* proteome is more rhythmic under entrained than constant conditions is consistent with previous observations in mouse^15^ and the overall percentage of clock-regulated proteins is within the range observed with other eukaryotes^13, 14, 21^. Rhythmic proteins in LL exhibited a mean period of oscillation of 23.3 h (Supp. Fig. 1A), in line with previous observations with clock luciferase reporter lines under similar conditions^22^.

**Figure 1:**
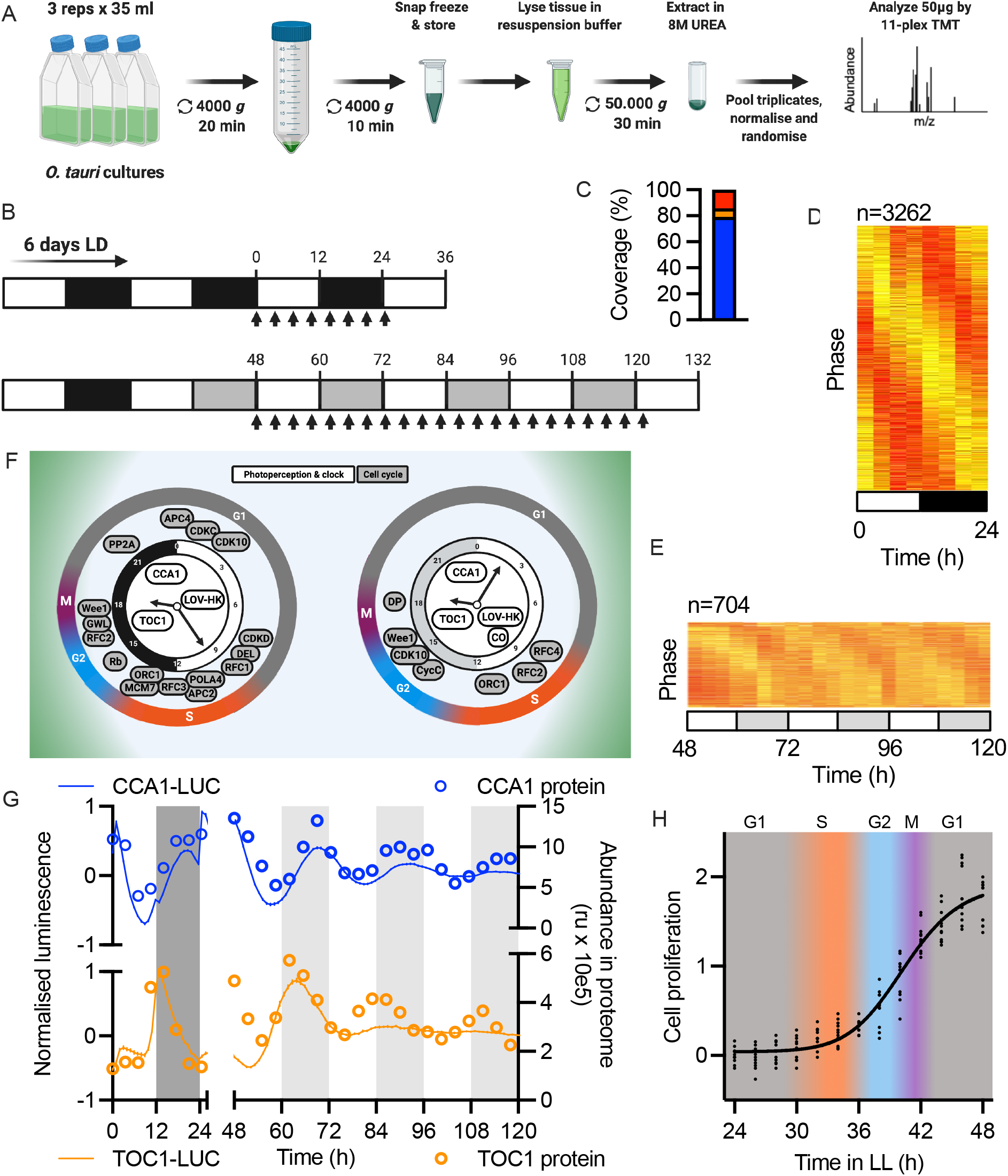
Deep-coverage diurnal and circadian proteomes. An overview of the experimental workflow (A) and sampling schedule (B) to obtain a deep-coverage proteome. Samples were taken every 3.5 hours (arrows) for 1 day under entrained conditions (LD) and for 3 days under constant conditions (LL). C) The percentage of nuclear-encoded *Ostreococcus* proteins that were quantified at all time points (blue), some time points (orange) or none (red). D-E) Heat maps showing min-max (red-yellow) normalised plots for all rhythmic proteins in LD (D) or LL (E). Rows represent individual proteins, ordered by phase, where each column is a separate time point. F) Diagrams depicting the key rhythmically abundant proteins of the circadian TTFL and photoperception (white) or the cell cycle (grey) expressed on a 24h clock face based on their peak phase under LD (leR) or LL (right). G) The relative abundance of TOC1 and CCA1 in LD (0-24 hours) and LL (48-120 hours) as determined by proteomics (right Y-axis; lines with data points) or luciferase results (leR Y-axis; mean with SEM). H) The cell cycle phases inferred from proteomics, overlaid with observed cell division events under LL.

Consistent with previous reports^20, 23-25^, oscillations in the abundance of the central components of the canonical TTFL circuit and light perception were phased similarly under the two conditions (Fig. 1F and Supp. Fig 1B-C). We compared the protein profiles of the *Ostreococcus* clock proteins CCA1 and TOC1^20, 26^ with longitudinal imaging of translational fusions of these proteins with firefly luciferase. We observed excellent agreement of phase and amplitude under LD, and a reduction in the dampening of oscillations under LL compared to what is commonly observed with luminescent marker lines in a sealed microplate (Fig. 1G).

As in other species^27^, the circadian clock and cell cycle are tightly coupled in *Ostreococcus* and show identical periods^28^, leading to the prediction that cell cycle proteins would show similarly organised oscillations between both conditions. We identified 44 core cell cycle candidate proteins based on the *Ostreococcus* genome (Supplemental File 1). Whilst more of these proteins showed significant daily rhythms under LD than LL, the cell cycle stages inferred from these rhythmically abundant cell cycle proteins are near-identical (Fig. 1F). Our data provide an unprecedented insight into the coupling of the cell and circadian cycles, indicating that progression into G1, S, G2, and M phases is initiated by a handful of regulatory proteins (Supp. Fig. 1D-F). As independent verification of inferred cell cycle stages, we monitored cell division under constant conditions and observed the anticipated relationship (Fig. 1H). The clock protein luminescence data (Fig. 1G) and matching cell division phases (Fig 1H) validate the integrity of our proteome dataset as an accurate description of temporal cellular organisation.

### Limited relationship between entrained and free-running rhythms

The consistent phase of the circadian and cell cycles between LD and LL conditions suggested that clock-regulated output pathways may also be similarly phased. Photosynthesis is one of the most important clock-regulated outputs, but when comparing the rhythmicity parameters of photosynthetic proteins there was little similarity or consistency between the identity or phase of rhythmically abundant proteins under LD compared with LL (Supp. Fig. 2). While this might be surprising at first sight, the divergence between LD and LL rhythms is very clear on the proteome-wide scale. Firstly, we observed limited overlap between rhythmic proteins under LD and LL conditions, indicating a complex interaction between endogenous circadian rhythms with environmental inputs (Fig. 2A). We also observed a significantly higher relative amplitude of rhythmic proteins under LD conditions (Fig. 2B), and although the overall difference in means is small, the maximum relative amplitude under LD was 94% versus 38% under LL. Even more notably, the phase distributions of rhythmic proteins were highly differential between LD and LL. The single predominant phase of highest abundance observed under LD was shortly before dawn, while under LL conditions this is in the early subjective night (Fig. 2C). While this difference may be explained in part by the small overlap between proteins rhythmic in LL and LD, the peak abundance phase is distinct even among those proteins that are rhythmically abundant under both conditions (Fig. 2D). Together, these data show a limited correlation in the identity of rhythmic proteins as well as their phase of peak abundance between circadian and diurnal conditions. This means that rhythmicity under entrainment cannot be inferred from circadian studies under constant conditions alone. As expected, there was no coherence between overrepresented Gene Ontology (GO) or Kyoto Encyclopedia of Genes and Genomes (KEGG) pathways at different phases under both conditions (Supp. Fig. 3). Therefore, the similarly phased cell and circadian cycles do not lead to coherent functional regulation of the proteome under LD versus LL conditions.

**Figure 2:**
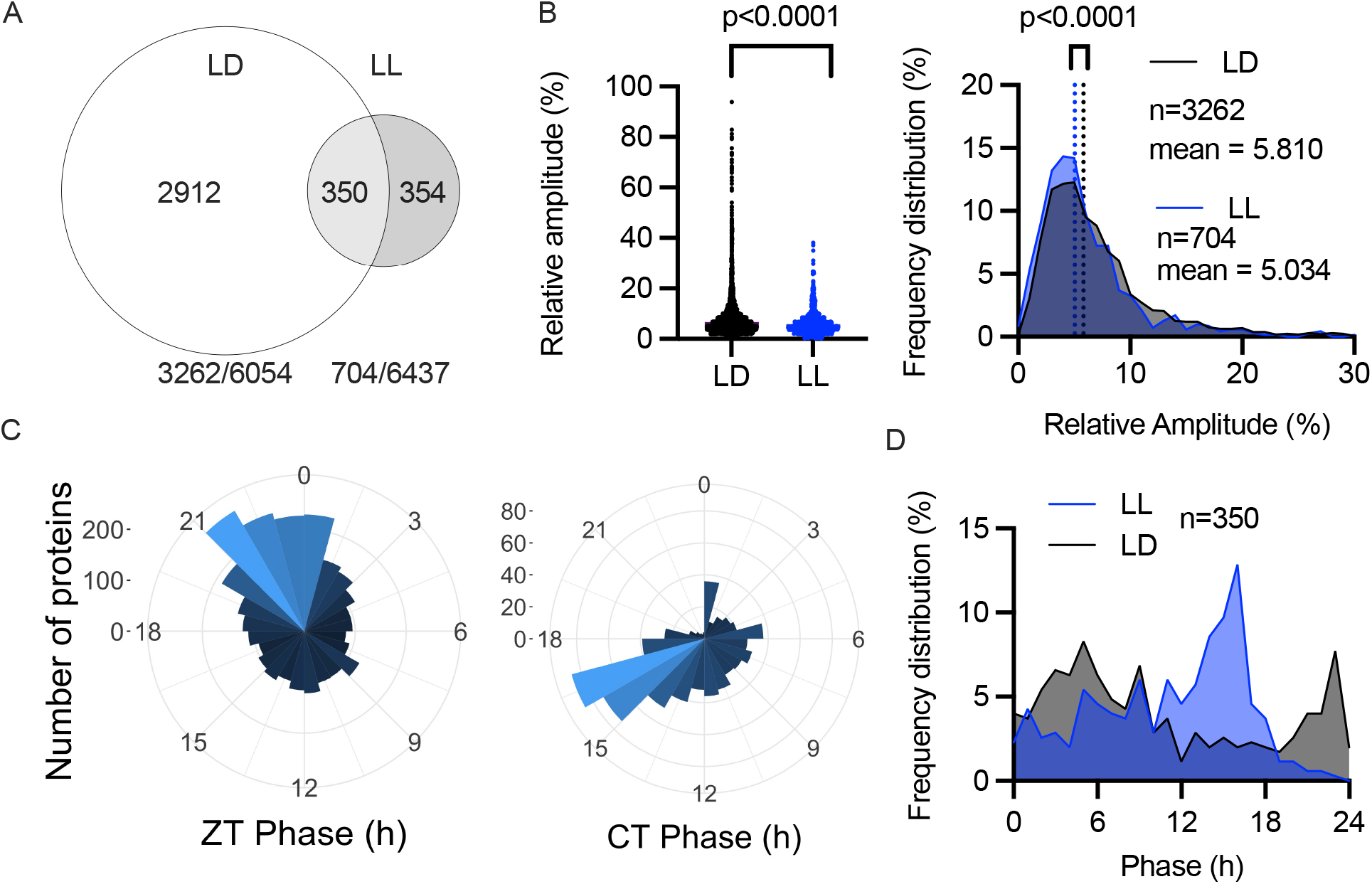
No clear relationship exists between entrained and free-running temporal proteomes. A) Venn diagram showing the overlap between rhythmic proteins under LL and LD, along with rhythmicity percentage under each condition. B) The relative amplitude of rhythmic proteins under LD or LL, expressed individually (leC) or as frequency distributions (right). C) Circular histograms showing the number of rhythmic proteins in LD (leC) and LL (right) at each 1-hour peak phase interval. D) Phase distribution under LD or LL of those proteins significantly rhythmic in both. Statistics reflect Mann-Whitney tests.

### Limited correlation between transcript and protein abundance rhythms

Transcriptomic data in *Ostreococcus* is currently limited to a single microarray study, sampled under entrained LD conditions^29^. The gene models used for that study^30^ have more recently been updated^31^, and upon re-assessment of the probe sequences, we found that 5925 out of 8056 probes map to a unique mRNA. We subjected the microarray data for these probes to the same rhythmicity tests as our protein dataset and found that nearly all transcripts (98%) exhibited significant rhythms in abundance under diurnal conditions (Fig. 3A), even though we found that only half of that number of proteins are rhythmically abundant under identical conditions. The peak phase distribution of transcript rhythms reveals an astonishing bimodal distribution, with peaks in phase around ZT9 and ZT20 (Fig. 3B) just before dusk and dawn at ZT12 and ZT0, respectively. This previously undetected double wave of transcriptional regulation is consistent with the canonical model of circadian rhythmicity, in which TTFLs allow cells to prepare for the differential demands of day and night through anticipatory changes in gene expression. Implicit to this model is the assumption that mRNA abundance determines protein abundance. However, we observed a stark contrast between transcript profiles and rhythmic protein abundances, as the latter shows only a single peak during 24h (Fig. 2C), in the late night/early morning.

**Figure 3:**
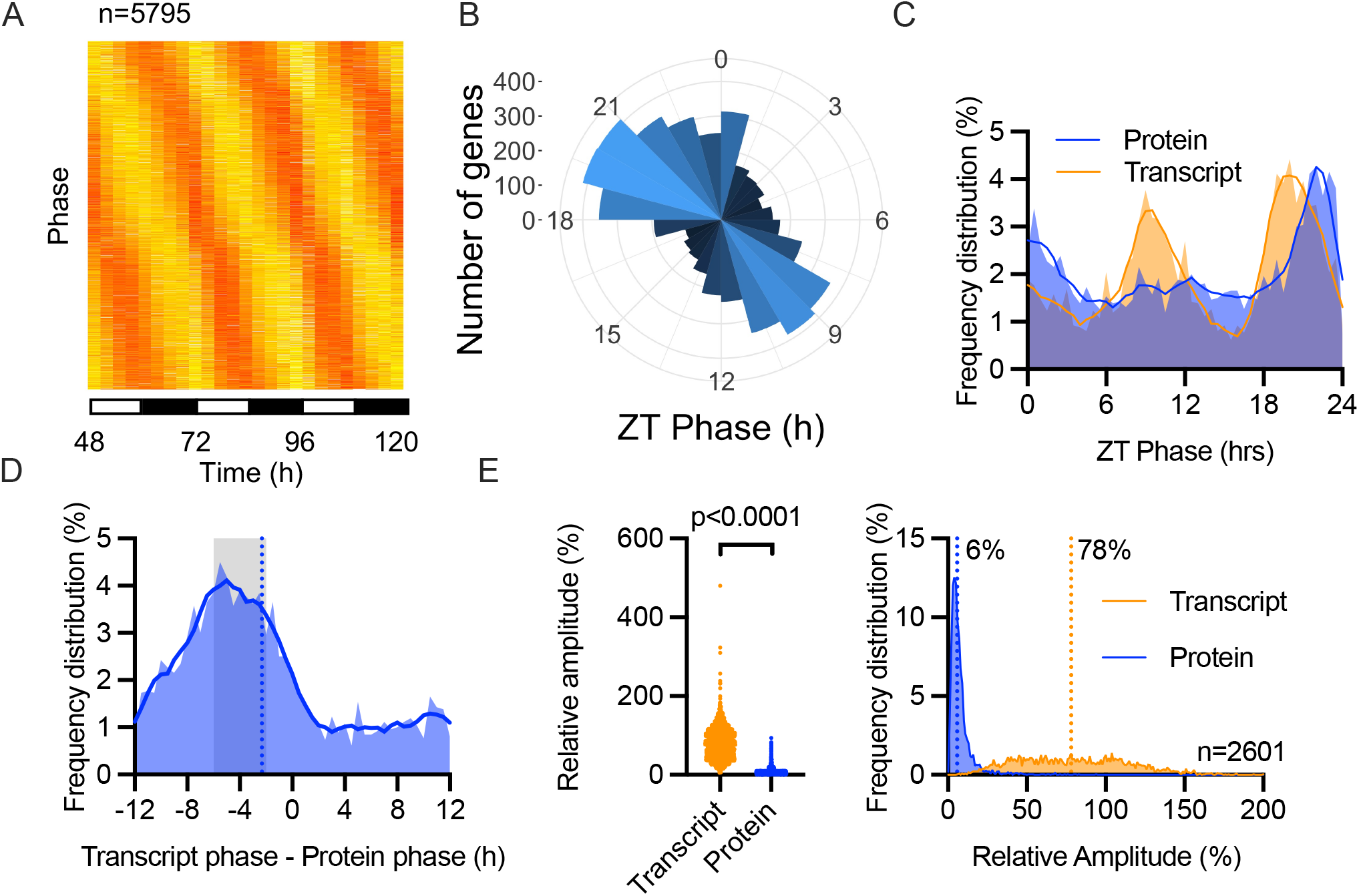
Partial correlation between transcript versus protein abundance. A)Heat maps showing min-max (red-yellow) normalised plots for all rhythmic transcripts following a reappraisal of published data^29^ under entrained conditions. B) Circular histogram showing the number of rhythmic transcripts in LD at each 1-hour peak phase interval. C) Frequency distribution of protein or transcript peak phase under LD conditions, for those where both gene products were rhythmic. D) One-for-one phase harmonics between rhythmic proteins and their rhythmic transcript, expressed as peak phase of transcript minus peak phase of protein. The mean value is indicated by a doLed line, and values that correlate with a protein peaking in a 2-6 hour window aNer the transcript are indicated. E) Relative amplitude values (leN) and frequency distribution of relative amplitude (right) of transcripts versus proteins under LD conditions. Statistics reflect Mann-Whitney tests.

To resolve this apparent paradox, we examined the temporal relationship between those rhythmically abundant proteins that are encoded by rhythmically abundant mRNAs, since it seemed plausible that the transcripts peaking at ZT9 might not encode rhythmic proteins. We identified 2601 such genes among the 4896 genes for which both transcript and protein abundance data is available. Again, we observed 2 daily peaks of transcript abundance but only one for protein peak (Fig. 3C). In a gene-by-gene comparison of the phase relationship between each transcript with its encoded protein (transcript phase – protein phase), the modal group of proteins peaked 4 hours later than their transcripts (Fig. 3D). Clear harmony was observed for the transcript peak preceding the protein peak of the canonical clock protein TOC1 and the photoreceptor LOV-HK, likely involved in entrainment (Supp. Fig. 4A).

These observations are consistent with findings from other organisms^13, 32^, and at first glance they support the canonical model of linear information flow in genetic systems (gene>mRNA>protein). However, only about 40% of the proteins peaked between 2 to 6 hours after their transcript, meaning that the majority of proteins did not (Fig. 3D). For example, the transcript and protein peaks of the key cell cycle kinase Wee1 were nearly antiphasic to one another (Supp. Fig. 4B). This corresponds with observations in other organisms that fewer than half of rhythmic proteins are encoded by rhythmic mRNAs under entrained conditions^15, 33^. Furthermore, transcriptional rhythms showed an average relative amplitude of 78%, with transcript levels oscillating by as much as 5-fold (Fig. 3E). This contrasts with the mean relative amplitude of 5.7% observed for protein levels under the same conditions, with no proteins exceeding a 1-fold change. The most parsimonious explanation is that the combined effects of the light-dark cycle and circadian system exert an effect on proteostasis at the post-transcriptional and post-translational levels that interacts with and modulates rhythmic transcriptional information flow from genome to proteome.

### Properties of rhythmic versus arrhythmic proteins

Considering the limited correlation between transcript and protein rhythmicity, we hypothesised that some of the variation we observe between rhythmically abundant proteins may be attributable to differences in their biochemical properties, which may confer a propensity or recalcitrance to circadian regulation. We therefore compared the properties of rhythmic versus arrhythmic proteins under LD and LL (Supp. Fig. 5A,B), but no significant differences were observed in protein isoelectric point, for example. However, we did notice that under either condition rhythmic proteins were significantly more likely to contain more predicted disordered regions than arrhythmic proteins. This could be linked to the observation that some clock proteins contain regions of disorder that are essential for their correct functioning^34 35^. We also found that protein size was significantly different between rhythmic and arrhythmic proteins under both conditions. While under LD, rhythmic proteins tend to be larger than arrhythmic proteins, the opposite was observed under LL. Under either condition, arrhythmic proteins were more hydrophobic than rhythmic proteins. However, the overall proteome-wide effect size of these differences in protein biochemical properties was modest and it seems unlikely that any of the factors we considered are sufficient to account for whether or not each protein’s abundance changes over the daily cycle.

**Figure 5:**
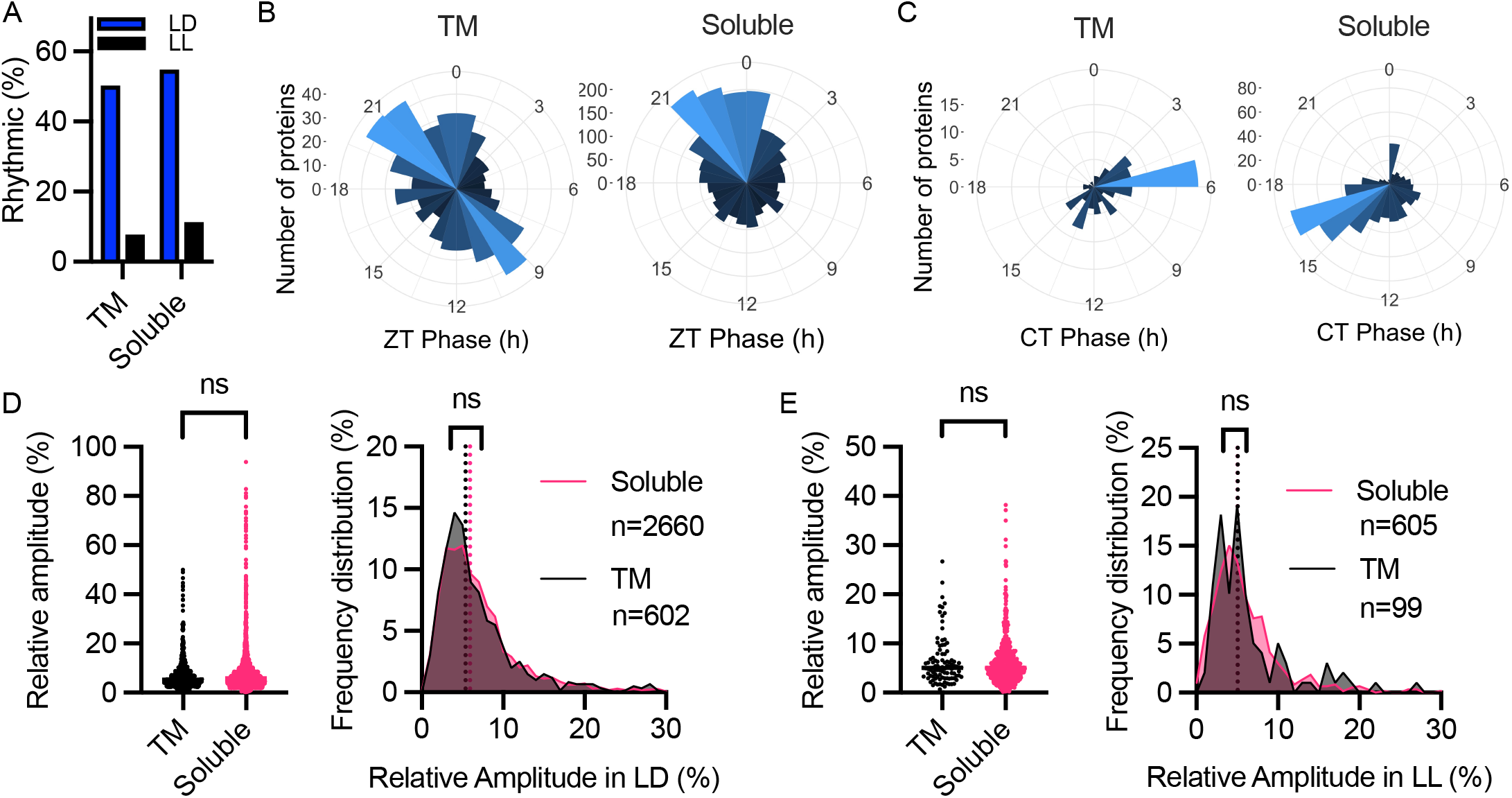
Proteins with transmembrane helices are differen8ally regulated from soluble proteins. A) Bar graph showing that proteins with transmembrane helices show a lower overall level of rhythmicity than those that do not, both under LD (blue) and LL (black). B-C) Circular histograms showing the number of rhythmic transmembrane (leB) or soluble (right) proteins in LD (B) or LL (C) at each 1-hour peak phase interval. D-E) The relaFve amplitude of transmembrane versus soluble proteins in LD (D) and LL (E) expressed individually (leB) or as frequency distribuFons (right). StaFsFcs reflect Mann-Whitney tests (ns = p>0.05).

We next considered whether the overall abundance of each protein might account for some of their rhythmic abundance under LD or LL. The overall mean abundance of all detected proteins under LD and LL was identical (dashed lines, Fig. 4A), indicating that overall cellular proteostasis is maintained between the two conditions. However, rhythmic proteins were significantly more abundant than arrhythmic proteins under entrained conditions (Fig. 4A, left), consistent with observations in mouse primary fibroblasts^36^. Interestingly, the opposite is true under LL conditions: rhythmic proteins were significantly less abundant than arrhythmic proteins (Fig. 4A, right). Despite these overall trends, the 660 proteins that were rhythmic in LL were more abundant than they were in LD (Fig. 4B and Supp. Fig. 5C,D). This effect was observed regardless of whether proteins were rhythmic in LD (310 proteins, 10% increase) or not (350 proteins, 6% increase). The 5394 proteins that were arrhythmic in LL did not significantly change in abundance compared to LD. Taken together, an increase in abundance from LD is associated with clock-regulated abundance rhythms in LL. However, similar to our analysis of mRNA rhythmicity and protein biochemical properties, protein abundance alone was not a reliable predictor of protein rhythmicity.

**Figure 4:**
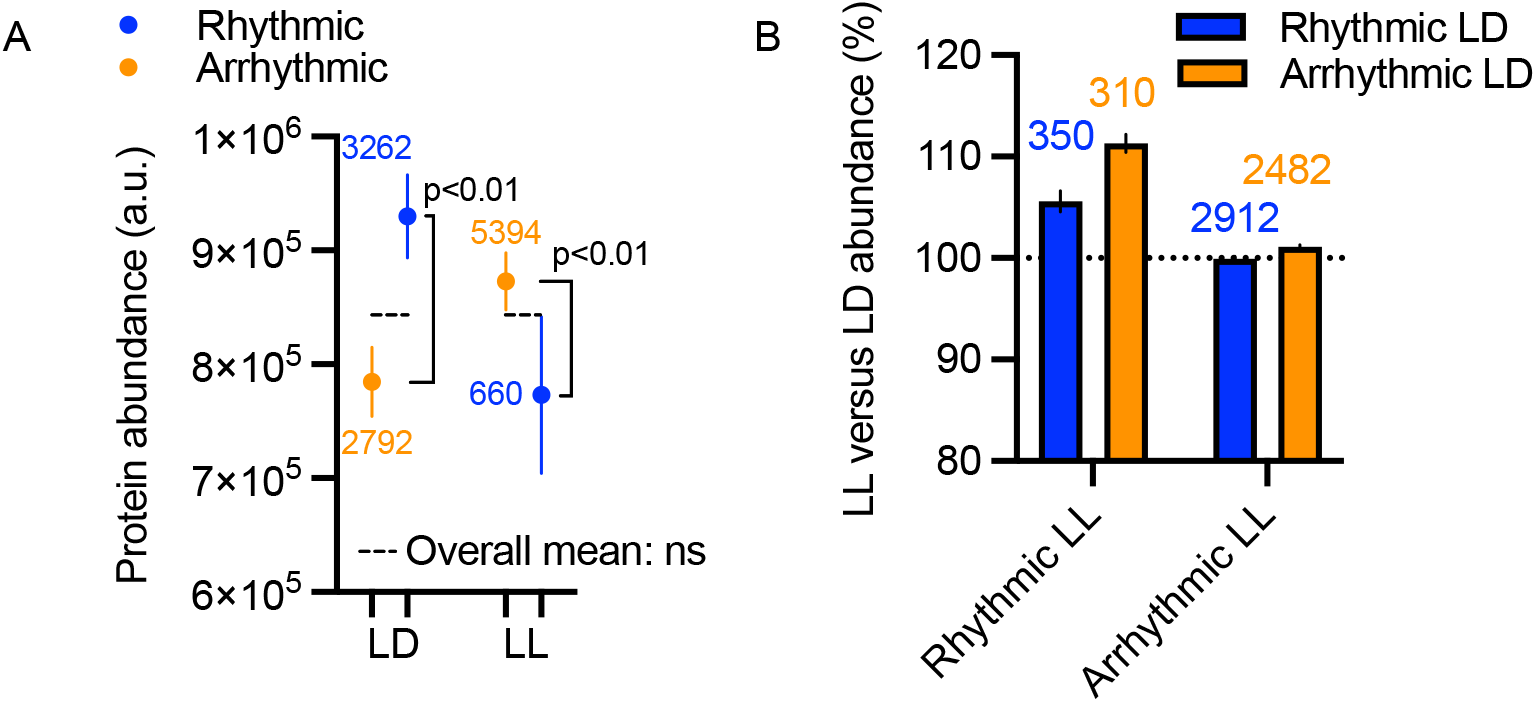
The differences between the abundance of rhythmic versus arrhythmic proteins is opposite under entrained or constant condi;ons. Mean protein abundance under LD and LL for rhythmic (blue) and arrhythmic proteins (orange). Dashed lines indicate the mean abundance under each condition. Number of proteins in each of the four groups are indicated. Statistics reflect Mann-Whitney tests (ns = p>0.05), error bars are SEM. B) Abundance in LL over LD, for proteins that are either rhythmic or arrhythmic in LL, separated by (ar)rhythmicity in LD. Error bars are SEM.

### Differential proteostasis of transmembrane and soluble proteins

Prompted by the observation of lower hydrophobicity in rhythmic proteins (Supp. Fig. 5A,B), we compared proteins with predicted transmembrane helices (TM, 1423 proteins) to those without (Soluble, 6276 proteins). A higher proportion of soluble proteins were rhythmic compared with TM proteins under LD (55% vs. 50%) as well as LL (11% vs. 8%); Fig. 5A). This observation contrasts with comparisons in the mouse liver proteome, in which transmembrane proteins were more rhythmic than soluble proteins^13^. Interestingly, the peak phase distribution of rhythmic transmembrane proteins was significantly different from rhythmic soluble proteins: in LD, the phase distribution of TM proteins was bimodal with peaks late in the day and late in the night, whereas soluble proteins peak only at the latter phase (Fig. 5B). Under LL conditions, TM proteins peaked in the middle of the subjective day, whereas soluble proteins peak during the subjective night (Fig. 5C). Peak phases of soluble proteins match the overall proteome peak phases, consistent with soluble proteins dominating the total proteome, while TM protein phases clearly deviate. The mean relative amplitude of rhythmic TM and soluble proteins was not different, but the maximum relative amplitude of soluble proteins was almost double that of TM proteins in LD as well as LL conditions (Fig. 5D). These results reveal a lower level of abundance rhythms among TM versus soluble proteins. Conversely, TM proteins were more heavily phosphorylated and more rhythmically phosphorylated than soluble proteins (Supp. Fig. 6). The highly conserved clock kinases CK1, CK2, and GSK3 are involved in circadian regulation in *Ostreococcus*^6, 37, 38^, yet they were not differentially abundant. These results would correlate with previous studies in mammals showing rhythmic phosphorylation state but constant protein levels for clock-related kinases^39, 40^. Together, we conclude that transmembrane proteins are less likely to be rhythmically abundant than soluble proteins, but are more subject to post-translational regulation, which might facilitate circadian regulation of transmembrane transport activity. Additionally, the phase separation between TM and soluble proteins suggests a rhythm in membrane protein biogenesis or degradation that occurs by a previously undetected and completely different process to soluble proteins.

**Figure 6:**
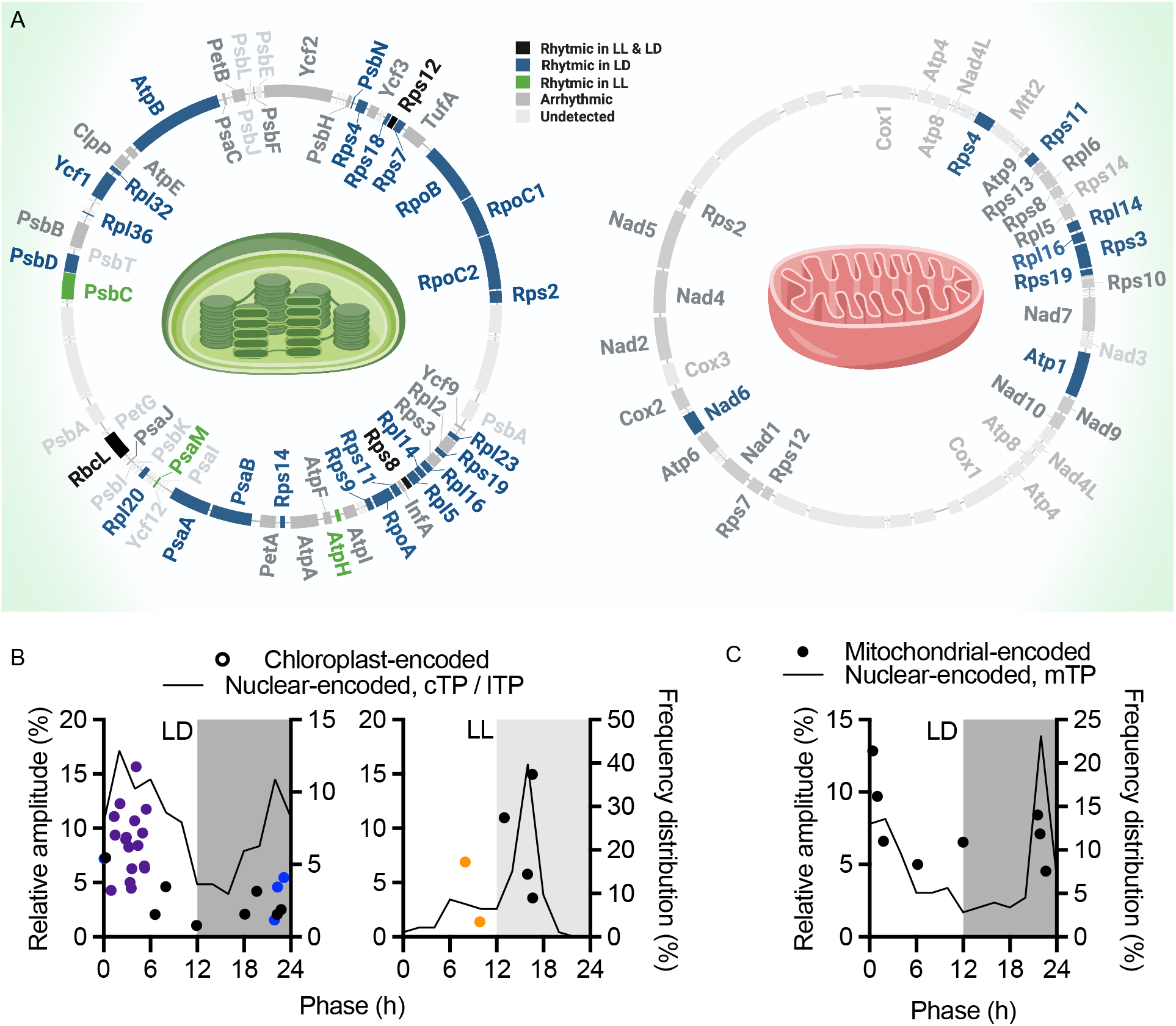
Orchestrated rhythmicity of organelle-encoded and organelle-targeted proteomes. Overview of the plas2d (le5) and mitochondrial (right) genome, with rhythmic, arrhythmic and undetected proteins annotated. B) Phase versus amplitude (le5 Y-axes) of rhythmic chloroplast-encoded proteins (data points, le5 Y axes) under diurnal or constant condi2ons. Purple dots represent ribosomal proteins, blue RNA polymerases, and orange photosystem components. Overlaid is the frequency distribu2on (black line, right Y-axes) of peak phase for rhythmic nuclear-encoded proteins carrying a chloroplast Transit Pep2de (cTP) or thylakoid lumen Transit Pep2de (lTP). C) Phase versus amplitude of rhythmic mitochondrial-encoded proteins (data points, le5 Y-axis) under diurnal condi2ons, overlaid with the frequency distribu2on (black line, right Y-axis) of nuclear-encoded proteins carrying a mitochondrial Transit Pep2de (mTP).

### Coordinated regulation of the organellar proteomes

Following the surprising phase separation of TM versus soluble proteins, we next analysed the organellar proteomes to deepen our understanding of spatial proteome regulation. *Ostreococcus* cells contain a single chloroplast and a single mitochondrion, containing autonomous genomes of 72 and 44 kb, respectively^41^. We detected 79% of the chloroplast-encoded and 63% of the mitochondrial-encoded proteome (Supplemental File 1). Under LD, 48% and 26% of the detected chloroplast- and mitochondrial-encoded proteins were rhythmic (Fig. 6A). The majority of the rhythmic chloroplast-encoded proteins were ribosomal proteins, which peaked in the first half of the day following the two chloroplast-encoded RNA polymerases that peak just before dawn (Fig. 6B, left). A smaller number of chloroplast-encoded proteins were rhythmic under LL (Fig. 6B, right)). In contrast, none of the mitochondrial-encoded proteins were rhythmic in LL, indicating that the circadian system does not control mitochondrial gene expression and that any rhythms under entrained conditions must result from environmental inputs. Indeed, the majority of rhythmic mitochondrial-encoded proteins under entrained conditions peaked around dawn (Fig. 6C).

As the majority of the organellar proteome is made up of nuclear-encoded proteins that are translocated, we compared the organelle-encoded proteins with nuclear-encoded proteins carrying a signal peptide targeting it for chloroplast or mitochondrial localisation. The peak abundance phase of proteins with a chloroplast Target Peptide (cTP) or thylakoid lumen Target Peptide (lTP) was highly consistent with the phase of chloroplast-encoded proteins under LD as well as LL (Fig. 6B). Nuclear-encoded sigma factors are required for transcription initiation in the chloroplast and have been shown to confer timing information to the chloroplast in Arabidopsis^42^. We observed a rhythmically abundant sigma factor (SIG6) phased at ZT2 under LD conditions and CT10 under LL conditions that potentially mediates the environmental and circadian control over chloroplast gene expression in *Ostreococcus* (Supp. Fig. 7). Like mitochondrial-encoded proteins, nuclear-encoded proteins carrying a mitochondrial Transit Peptide (mTP) peaked around dawn under diurnal conditions (Fig. 6C). Overall, the striking similarity between organelle-encoded and organelle-targeted protein peak abundance phases indicates a purposeful and coordinated temporal regulation of chloroplast and mitochondrial function. Our cell cycle analysis (Fig. 1) suggests that this observed peak phase for organellar proteins is consistent with the time of organelle duplication, ahead of cell division later in the night.

## Discussion

We explored the spatiotemporal regulation of a eukaryotic cellular proteome in unprecedented depth. This revealed a modest overlap between the rhythmic proteomes under LD and LL, which indicates complex interactions between clock-regulated rhythms and daily environmental cycles, with a larger number of proteins regulated by the latter. We found there is a little consistency between transcript and protein rhythmicity, and identified a previously unknown disconnect between the peak abundance phases of transmembrane and soluble proteins. Conversely, the peak abundance phases of organelle-encoded and organelle-targeted proteins show a high degree of synchrony. We also observed small differences between the biochemical properties of rhythmic versus arrhythmic proteins. However, the key observation that permeated through all our analyses was that no single variable was able to account for protein abundance rhythms.

Therefore, rhythmic regulation of protein abundance most likely involves both specific as well as more general mechanisms that together determine the relative balance between protein synthesis versus turnover of each protein. Certainly the global rates of transcription, translation, and protein degradation are subject to circadian regulation^9, 43-46^. Additionally, transcriptional, post-transcriptional and post-translational regulation is often light-dependent^15, 29, 47, 48^. For example, while the degradation rate of *Ostreococcus* CCA1 protein is rhythmically regulated by the circadian clock, the degradation rate of TOC1 is instead regulated by dark-to-light transition^45^. Together, these novel considerations ask for a reappraisal of the fundamental bases of cellular rhythms.

The canonical circadian model suggests that circadian regulation of cell function is driven solely by rhythmic gene expression, established by transcriptional/translational feedback in the ‘core clock’. This model assumes linear flow of information: rhythmic regulation of gene expression leads to rhythmic mRNAs, which leads to rhythmic protein levels and rhythmic function. The evidence for rhythmic regulation of transcript abundance and the role of clock proteins within that is irrefutable. However, the poor correlation we observed between transcript and protein rhythmicity parameters argues that rhythmic transcription simply cannot be assumed to elicit comparable (or indeed any) changes in protein activity. Evidence from other eukaryotes supports the notion that rhythmicity of protein abundance cannot be inferred from rhythms in transcript abundance^33, 46, 49, 50^. This is consistent with a growing number of studies from outside the circadian context that prove that mRNA abundance is poorly predictive of protein abundance or activity^51-53^. In addition to the large body of evidence for the existence of circadian timekeeping without daily cycles of transcriptional activation/repression^54^, the paradigm that transcript rhythmicity is the only physiologically relevant mechanism of timekeeping is no longer tenable.

A second model that might better account for circadian cellular rhythms, known as ‘revised canonical’, assumes that rhythmic translation of existing mRNA leads to proteome-wide rhythms, which in turn drive rhythmic function. In this model, any mRNA rhythms would be of secondary importance so long as sufficient template is present for each protein’s translation. There is sufficient evidence to support that global translation rates are indeed rhythmically regulated^9, 44^. However, this model would predict similar peak abundance phases and a far higher overlap between the identity of rhythmic proteins under entrained and constant conditions. Additionally, it cannot explain the different phase of transmembrane versus soluble proteins, nor the coordination of organelle-encoded with organelle-targeted proteins. Therefore, we ultimately find no evidence to support this model.

A third model for circadian rhythms is that a post-translational oscillator drives rhythmic cell function through rhythmic protein modification and/or degradation rhythms. Here rhythms in transcript abundance result from rhythmic protein activity instead of the other way round, i.e. rhythms in any mRNA or protein abundance are not the cause but a consequence of rhythmic post-translational activities. Prior evidence in *Ostreococcus* for rhythmic regulation in the presence or absence of rhythmic gene expression includes redox-sensitive post-translational modification rhythms and rhythms of Mg^2+^ and K^+^ transport over the plasma membrane. The observed enrichment for transmembrane transporters among rhythmically phosphorylated proteins likely accounts for these previously reported ion transport rhythms, which is underlined by the low prevalence of transmembrane transport proteins among rhythmic proteins. Our observations that the differential abundance of rhythmic and arrhythmic proteins is opposite under entrained versus constant conditions, and that an increase in mean abundance upon transition from LD into LL is associated with rhythmicity, are only consistent with this third model and not the first two.

Therefore, our study provides support for the idea that a combination of transcriptional and post-translational regulation accounts for how ∼24h regulation of eukaryotic cell biology is achieved, similarly to the cyanobacterial circadian system^55^. On one hand it is clear from the observed phase harmonics that transcriptional rhythms exert some level of control over the proteome. On the other hand, the small relative amplitude of rhythmic proteins (∼5%) certainly begs the question of whether protein abundance rhythms are in fact a functionally relevant output of transcriptional rhythms. It seems more likely that transcript rhythmicity presents a means of achieving proteostasis, to counteract a rhythmic requirement driven by rhythmic function and associated rhythmic protein turnover. This interpretation would be supported by the recent observation in mammalian cells that more rhythmic proteins are observed without a transcriptional ‘core clock’ than there are observed with one^36^. Therefore, we suggest that perhaps the canonical transcriptional/translational clock model might be back to front: it is more consistent with available data that rhythmic function generates rhythmic gene expression as a consequence of rhythmic post-translational regulation and protein degradation.

Overall, this blueprint for the temporal proteomic landscape of a eukaryotic cell provides valuable lessons on the information flow underlying cellular rhythms. Ultimately, to fully understand eukaryotic cellular timekeeping, a complete picture of circadian post-transcriptional, translational, and post-translational regulation is needed alongside transcriptional regulation. To obtain comprehensive models of the cellular circadian landscape, it will be necessary to integrate different levels of organisation by multi-omics approaches. This integration relies on high-quality datasets, and by contributing the most detailed temporally resolved proteome across Eukaryota, we have provided a significant step in this direction.

## Methods

### Sample collection and preparation

*Ostreococcus tauri* cells were cultured in artificial seawater (ASW) and entrained under cycles of 12 h light: 12 h dark (LD) as reported previously^9^ for 6 days. 24h prior to the first sampling point, cultures were either transferred to constant light (LL) or kept in LD cycles (LD, Fig. 1A). Samples were collected in triplicate every 3.5 hours for 3 full cycles in LL (22 time points) and one full cycle in LD (8 time points). For each time point whole cells were collected at 4000 rpm for 20 min. Media was discarded and each cell pellet was gently resuspended in 0.9 ml ASW. Cells were collected at 4000 rpm for 10 min and the media discarded. After adding two chrome beads (3 mm) to each pellet the samples were snap-frozen in LN2. Samples were stored at −80 °C until all time points were collected. Resuspension buffer (50mM Hepes, pH7.5, 150mM NaCl, protease inhibitors (Roche)) were added to each frozen pellet on ice. Cells were lysed using a Tissue Lyser (Eppendorf) in precooled blocks (1 min at 30/s). Whole lysates were centrifuged at 50,000g for 30 min at 4°C in a Beckman Optima MAX ultracentrifuge. Pellets were kept on ice and were washed once carefully with resuspension buffer. Pellets were resuspended in 8 M urea buffer (8 M urea, 20 mM Tris-HCl, pH 8) by vortexing. Samples were then sonicated in a Bioruptor (Diagenode) for 30s on/ 30s off (x5). All samples were centrifuged at 17,000g for 10 min to remove unsolubilised debris. Triplicate samples per time point were pooled, samples were randomised, and 50 µg of each time point was analysed by 3 sets of 11-plex TMT.

### TMT peptide labelling

Samples were randomised before allocating to TMT runs, and the operator was blinded to the sample IDs. Samples were trypsin-digested as reported previously^36^. Lyophilized peptides were resuspended in 20 µl of 175 mM triethylammonium bicarbonate and labelled with a distinct TMT tag, 12µl, from a stock prepared as per manufacturer’s instructions (Thermo Scientific), for 60 minutes at room temperature. The labelling reaction was quenched by incubation with 2.2 µl 5% hydroxylamine for 30 min. Labelled peptides from 10 time point samples and 1 pool were combined into a single sample and partially dried to remove MeCN in a SpeedVac (Savant). Samples were desalted and the eluted peptides were lyophilized.

### Basic pH Reverse-Phase HPLC fractionation and LC-MS/MS

The TMT labelled peptides were subjected to off-line High-Performance Liquid Chromatography (HPLC) fractionation as described [wong et al, biorxiv]. The fractionated peptides were analysed by LC-MS/MS using a fully automated Ultimate 3000 RSLC nano System (Thermo Scientific) fitted with a 100 μm x 2 cm PepMap100 C18 nano trap column and a 75 μm × 25 cm reverse-phase NanoEase M/Z HSSC18 T3 column (Waters). Samples were separated using a binary gradient consisting of buffer A (2% MeCN, 0.1% formic acid) and buffer B (80% MeCN, 0.1% formic acid), and eluted at 300 nL/min with an acetonitrile gradient. The outlet of the nano column was directly interfaced via a nanospray ion source to a Q Exactive Plus mass spectrometer (Thermo Scientific). The mass spectrometer was operated in standard data-dependent mode, performing a MS full-scan in the m/z range of 380-1600, with a resolution of 70000. This was followed by MS2 acquisitions of the 15 most intense ions with a resolution of 35000 and Normalised Collision Energy (NCE) of 33%. MS target values of 3e6 and MS2 target values of 1e5 were used. The isolation window of precursor ion was set at 0.7 Da and sequenced peptides were excluded for 40 seconds.

### Spectral processing and protein identification

Raw files were processed using MaxQuant^56^ v 1.6.6.0. MS/MS spectra were quantified with reporter ion MS2 and searched against the nuclear-encoded proteome obtained from the Ostta V2.2 database^31, 57^ plus the mitochondrial and chloroplastic genomes^41^. Carbamidomethylation of cysteines was set as fixed modification, while methionine oxidation, N-terminal acetylation and phosphorylation of serine, threonine and tyrosine, were set as variable modifications. Protein quantification requirements were set at 1 unique and razor peptide. In the identification tab, second peptides and match between runs were not selected. Other parameters in MaxQuant were set to default values. The MaxQuant output file was processed with Perseus (v1.6.6.0). Identifications from the reverse database were removed, only identified by site, potential contaminants were removed, and we only considered proteins with ≥1 unique and razor peptide. All columns with an intensity “less or equal to zero” were converted to “NAN” and exported as a .txt file. The MaxQuant output file with phosphor (STY) sites table was also processed with Perseus software (v1.6.6.0). The data was filtered: identifications from the reverse database were removed, potential contaminants were removed and we only considered phosphopeptides with localisation probability ≥ 0.75. Then all columns with intensity “less or equal to zero” were converted to “NAN” and exported as .txt file. The mass spectrometry proteomics data have been deposited to the ProteomeXchange Consortium via the PRIDE^58^ partner repository with the dataset identifier PXD025009.

### Normalisation

Since an equal amount of protein was used for each TMT labelling reaction, sample loading normalisation was performed by taking the sum of all intensities for each time point, and normalising to the mean of these. Internal reference scaling (IRS)^59^ was then carried out to allow for comparisons between TMT experiments: the mean abundance for each protein in each of the three pools was calculated. The mean of these means was calculated and used to normalise the value for each protein between the three TMT runs. In instances where a protein was missing from a pool sample, the mean of the remaining 2 pool samples was used to normalise. For all except five proteins in the dataset, if a protein was missing from the pool sample it was also missing from every individual time point in that TMT run. Peptides that are not unique to one protein were removed, as were proteins that were detected at too low levels to reliably quantify. Time points were then de-randomised to obtain the final data set.

### Circadian parameter estimations

For all LL datasets, rhythmicity was tested using the JTK_cycle algorithm with empirical calculation of the p-values (eJTK)^60^ using BioDare2^61^. All time points were included in the rhythmicity analysis, with no detrending of input data, and parameters as preset for ‘eJTK Classic’. The cut-off for rhythmicity was at p<0.05, as customary for eJTK. As a single cycle of time series data is not sufficient for eJTK analysis^18, 62^, all LD data was assessed for rhythmicity using Cosinor analysis^63^ in GraphPad Prism, comparing a cosine wave with period constrained to 24h and a rate constant for signal decay to a horizontal line with a sum-of-squares F test. A cut-off of p<0.05 to prefer the cos wave was used to deem a protein significantly rhythmic. For all rhythmic proteins, BioDare2 was used to calculate all circadian parameters. The MFourFit algorithm was used for absolute phase and amplitude calculation with amplitude and baseline detrending. Data were log2 transformed prior to all rhythmicity and parameter calculations, except for amplitude values. LD data were concatenated to be able to approximate these parameters, and the period was constrained to 23.5-24.5 hours. The MESA algorithm with amplitude and baseline detrending was used for period calculation of the LL data, constrained to 18-34 hours. To transform absolute phase to circadian phase, absolute phase predictions by MFourFit were divided by 24, and multiplied by the period as estimated by MESA. Proteins with any missing values in the LD dataset were omitted from the analysis but kept in the data set with circadian parameters as Not Determined (ND). For the LL dataset, a maximum of 1/3rd of missing values was allowed, equating to those missing values for one out of three TMT runs. Phosphosite data were normalised as described above. For LL, phosphosites detected in two-thirds of time points were kept in the dataset, and for LD only those present in all time points were retained. Prior to rhythmicity and circadian parameter analyses, the phosphosite data were normalised to their protein abundance across the time points. Phosphosites for which the protein had not been detected in the dataset were removed. Rhythmicity and circadian analyses were carried out as above.

### Biological verification experiments

Luminescence experiments using CCA1-LUC and TOC1-LUC were performed as reported previously^22^. Raw luminescence data was detrended by subtracting a rolling average of luminescence readings for the following 24h. For cell proliferation analyses, cultures subjected to the identical conditions as described for the proteomic analyses were sampled every 2 hours on the second day of constant light. Cells were counted under a light microscope using a haemocytometer. Two biological replicates were performed, with 5 technical replicates for each time point.

### Transcript analysis

The probe sequences from a publicly available *O. tauri* microarray study under entrained LD conditions ^29^ were originally designed using outdated gene models and were therefore blasted against the *O. tauri* genome V2.2 on the Orcae service^31, 57^. This resulted in usable microarray data for 5925 of the 7700 genes in the genome. Circadian parameters were estimated as outlined for the LL proteome dataset above, using eJTK for rhythmicity analysis and MFourFit for phase and amplitude calculation, with period constrained to 23.5-24.5 hours.

### Structural and functional protein data

Gene ontology and KEGG pathway analyses were performed in R v3.6.1. The enrichGO and enrichKEGG functions from the clusterprofiler R package^64^ were used to compare a target dataset to background. Transmembrane helices in proteins were predicted using web tool TMHMM server 2.0^65^. Proteins with at least 1 predicted transmembrane helix were considered ‘transmembrane’ in subsequent analyses. The Sequence Manipulation Suite web tool was used to calculate protein molecular weight and isoelectric point^66^. N-terminal presequences in the entire nuclear-encoded proteome were identified with TargetP-2.0^67^. Those containing a predicted mitochondrial transit peptide (mTP) were used to generate a ‘mitochondrial proteome’, and those containing either a chloroplast transit peptide (cTP) or thylakoid luminal transit peptide (luTP) were used for the ‘chloroplast proteome’. Hydrophobicity and intrinsic disorder of proteins were calculated by localCIDER package^68^ in Python 3.x, with the get_kappa and get_uversky_hydropathy functions used for intrinsic disorder and hydrophobicity respectively^69, 70^. A low kappa value implies a propensity to form random coils and therefore higher intrinsic disorder. Radial plots were created using the ggplot2 R package. Venn diagrams were generated using the eulerr R package. To generate heat maps, the rhythmic proteins/transcripts were ordered by their calculated phase (absolute phase for LD proteins/transcript or phase in circadian time for LL proteins). The abundance of each protein was normalised by the time course mean of the protein, and values were centred around 0 using the scale function in R before applying the heatmap.2 function from the pvclust R package^71^. Relevant proteins from different biological processes were depicted in diagrams based on the following cell categories: cell cycle^28, 29^, light signalling and clock^25, 72, 73^, photosynthesis and chloroplast biosynthesis^74^, and organelle-encoded proteins^41^. Bioinformatic analyses to identify potential candidate proteins were conducted using Standard Protein Basic Local Alignment Search Tool (BLAST) of known protein sequences from other model systems as *Arabidopsis thaliana* and *Homo sapiens* using the *Ostreococcus* genome ORCAE V2^57^. Graphs, density plots and histograms were plotted and statistics were calculated using GraphPad Prism v8, R v3.6.1, or with BioRender.

## Supporting information

Supplemental Figures, Legends, and References

## Acknowledgements

The authors would like to acknowledge Jennifer Hurley, Jackie Pelham, Andrés Romanowski, Babette Vlieger, Francisco José Romero-Campero, Ana Belén Romero Losada, and Phil Kirk for bioinformatics support. EG was supported by a Wellcome Trust Institutional Strategic Support Fund award to GvO, HK by a Royal Society Research Fellows Enhancement Award to GvO (RGF\EA\180192), and SG by a Leverhulme Trust Research Grant awarded to GvO (RPG-2019-184). HF is a PhD student funded by the Biotechnology and Biological Sciences Research Council (BBSRC, BB/M010996/1). AS was supported by the AstraZeneca Blue Skies Initiative. JSO was supported by UKRI Medical Research Council (MC_UP_1201/4). GvO was supported by a Royal Society University Research Fellowship (UF110173) and renewal (UF160685).

## Author contributions

GvO and JSO conceived the approach. GvO provided the overall supervision and management of the project. The protocol development and all pilot studies were carried out by EG. Time series sampling and processing of samples were performed by EG with assistance from HF. AS contributed to developing the detection strategy using TMT labelling. SPC labelled the samples and performed mass spectrometric detection. Bioinformatic analysis of the resultant data and generation of figures was carried out by HK, SG, and GvO. HK, SG, and HF wrote the first draft. All authors interpreted the results. GvO wrote the final draft on which all authors commented.

## Competing interests

The authors have no competing interests to declare, financially or otherwise.

## Notes

### Competing Interest Statement

The authors have declared no competing interest.

