## Supplemental Figures, Legends, and References for "Differential effects of environmental and endogenous 24h rhythms within a deep-coverage spatiotemporal proteome"

*Supplemental Figure 1*

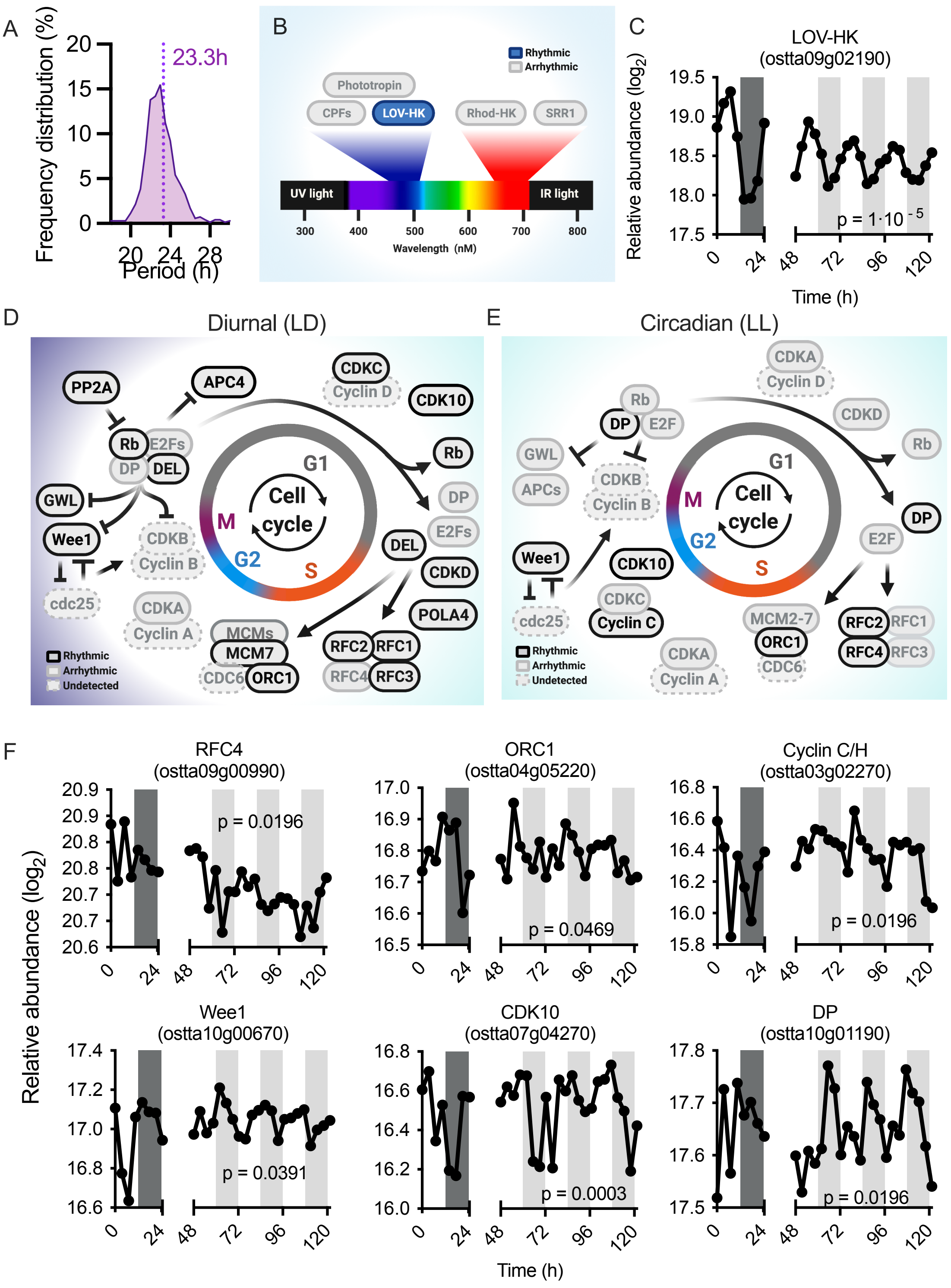


*For Legend, PTO*

*Supplemental Figure 1: Rhythmicity within the circadian clock and cell cycle*

A) A frequency histogram of the period of oscillations observed under LL shows the expected mean value of ~23h. B) Photoreceptors in the *Ostreococcus* proteome. *Ostreococcus* senses light changes in the environment through different blue and red light photoreceptors. The histidine kinase LOV-HK, Cryptochrome/Photolyase family (CPF) proteins, and Phototropin display UV/blue-light sensing properties, while red light fluencies are perceived by the histidine kinase Rhod-HK and SRR1[23]. We observed full coverage of all main photoreceptors*,* and remarkably rhythmic protein abundance is only evident for LOV-HK, under both entrained and free-running conditions. In *Ostreococcus*, LOV-HK is transcriptionally regulated by the clock around dawn and is also essential for synchronizing core clock components[24], not only upon blue light exposure but also in response to other light wavelengths, indicating interconnected light pathways. Interestingly, vertical distributions of *Ostreococcus* populations in aquatic environments suggest that UV/ blue light is the main light source due to the wide blue light distribution at depth, which would help cells to discriminate time of day[75, 76]. C) Traces of quantified data from LD and LL time series for the LOV-HK protein. P values refer to the eJTK score in LL. D-E) Reconstruction of the *Ostreococcus* cell cycle under LD (D) or LL (E), indicating the instructive rhythmic proteins. Our bioinformatic analyses identified 44 core cell cycle candidate genes in the *Ostreococcus* genome, of which we detected 35 of the protein products. Only 7 of these proteins were rhythmically abundant under LL and 16 proteins under LD. Interestingly, our analyses provide a detailed perspective on the tight control of cell cycle phases by the circadian clock under both conditions, with near-identical cell cycle stages. G1 proteins such as the CDKC and CDKD or E2F factors as DEL only displayed rhythmicity under LD, suggesting no direct role of the circadian clock modulating G1 phase protein abundance. Specific subunits of the S phase RFC complex and the Pre-replication complex were rhythmically abundant under both conditions (RFCs, MCMs and ORC1), indicating that coupling of the circadian clock to the progression through S phase is mainly mediated through the RFC and Pre-replication complexes. Although several studies show no discrete activity of Cyclin C at any specific cell cycle phase[77], we found that Cyclin C protein abundance under LL rhythmically peaks at CT15, matching with the G2 phase in which cells prepare for cell division. This result would correlate with the major role in RNA transcription initiation of Cyclin C and the key transcription event of M phase-related genes before cell division observed in humans[78]. While rhythmic protein abundance of the key kinase Wee1 peaked at a similar phase under both conditions, other kinases such as CDK10 displayed almost opposite rhythmic patterns under LL and LD. Wee1 is clock-regulated in other organisms and determines entry to M phase, and is also involved in other processes such as the DNA damage response[79-83]. Similarly, CDK10 plays important roles in genome stability and DNA repair. These results could indicate a circadian contribution to preventing genome instability and maintaining DNA integrity, by regulating the G2/M transition under LL through the protein levels of Wee1, and a more flexible regulation of CDK10 based on the light/dark cycles. During the M phase, although several proteins from the Retinoblastoma complex (Rb, DP, E2Fs as DEL) and phosphatases as PP2A showed clear rhythmicity under LD, DP protein was the only rhythmic protein under LL peaking in the middle of the night. DP protein is an essential component of the Retinoblastoma complex, which is assembled at the end of the M phase to start a new cycle by inhibiting M phase-related proteins[84-86]. This suggests a direct circadian regulation of the M phase through DP protein and additional light/dark regulation of other M phase proteins. F) Traces of quantified data from LD and LL time series for examples of cell cycle proteins significantly rhythmic under LL. P values refer to eJTK scores under LL. The raw data for all cell cycle proteins can be found in Supplemental File 1.

*Supplemental Figure 2*

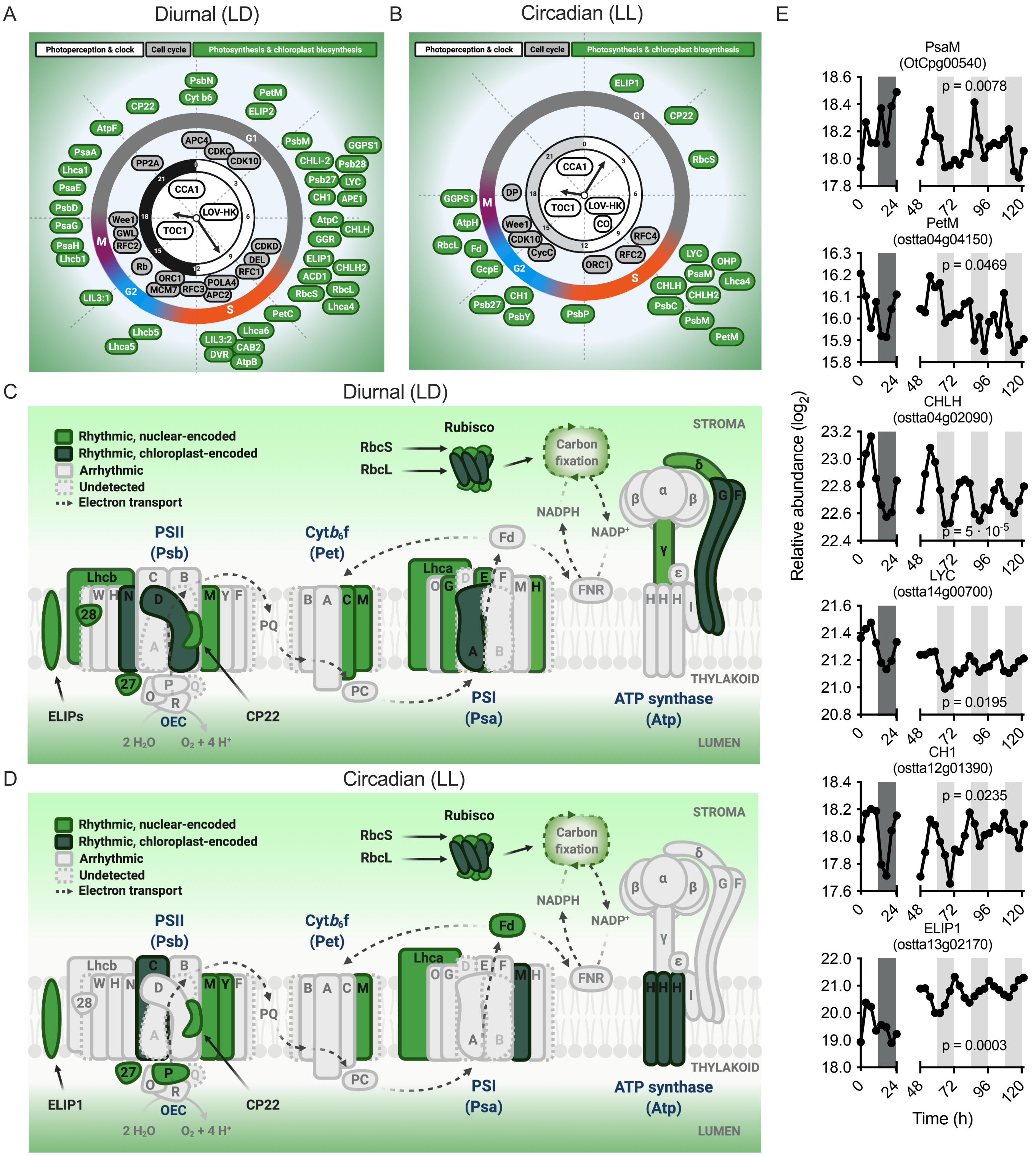


*For legends, PTO*

*Supplemental Figure 2: The rhythmically regulated proteins of the photosynthetic machinery*

A-B) Diagrams similar to Fig. 1F, but in addition to depicting the key rhythmically abundant proteins of the circadian clock and photoperception (white) or cell cycle (grey), also includes output rhythms in photosynthetic and chloroplast biogenesis-associated proteins expressed on a 24h clock face based on their peak phase under LD (A) or LL (B). C-D) Rhythmicity within the photosynthetic apparatus under LD (C) or LL (D) conditions. Circadian regulation of photosynthetic genes has been well described[42, 87, 88]. This apparatus consists of 4 main protein complexes, conserved from cyanobacteria to higher plants: Photosystem II (PSII), Cytochrome $b_6f$ (Cyt$b_6f$), Photosystem I (PSI), and ATP synthase. Among the 74 photosynthesis-related proteins we detected, a remarkably different organization of rhythmic proteins under LL compared to LD cycles was observed. All main photosynthetic complexes contain rhythmically abundant subunits. Specifically under LL, proteins from the photosynthetic apparatus (Psb, Pet, Psa) and pigment biosynthesis (Lycopene lyase (LYC) and Magnesium chelatases (CHLH, CHLH2)) were predominantly most abundant around CT8, coinciding with the start of the S phase of the cell cycle. Our results therefore match previous observations in algae[89] that suggest the onset of chloroplast division at the start of the S phase, ahead of cell cytokinesis. Consistent with that result, proteins involved in the later stages of photosystem assembly and chloroplast formation[74, 90], such as Psb27 and PsbY, peak subsequently. Under entrained conditions, abundance rhythms are more prevalent among photosynthetic proteins including those of the light harvesting complexes (LHCAs, LHCBs, LIL3 proteins), and these show a broader phase distribution over the light period. This indicates an important light/dark regulation of a subset of photosynthetic proteins separate from the circadian control. Proteins involved in energy-related processes at the end of the photosynthetic apparatus also showed differential rhythmicity under both conditions: subunits of the ATP synthase complex (AtpC, AtpF) were rhythmic under LD, but only the transmembrane ATPase subunit C (AtpH) and Ferredoxin peaked at night. Proteins involved in isoprenoid biosynthesis, such as Chlorophyll a oxygenase (CH1) and GeranylGeranyl Pyrophosphate Synthase 1 (GGPS1), peaked at opposite phases in LL compared to LD, which could correspond with known light-dark-dependent effects of downstream steps of pigment biosynthesis pathways[91-93]. Our results also showed rhythmic photoprotection-related proteins such as Early Light-Induced Proteins (ELIPs), CP22 or One Helix Protein (OHP) peaking during the day, which correlates with anticipation of photodamage by light exposure[94, 95]. Finally, the two subunits of the enzyme Rubisco (small subunit RbcS and large subunit RbcL) displayed differential rhythmic abundance between LD and LL. Subunits peaked at opposite phases under LL but at similar phases under light/dark cycles, supporting the notion that Rubisco subunits are translated and assembled in a time- and a light-dependent manner[96]. Overall, proteomics results integrate well with previous studies from the clock, cell cycle, and light signalling fields, and illustrate in detail how pervasive rhythmicity throughout cellular biochemistry is achieved by circadian rhythms in the abundance of specific components within pathways. E) Traces of quantified data from LD and LL time series for some examples of photosynthesis-related proteins significantly rhythmic under LL. P values refer to eJTK scores under LL. The raw data for all photosynthetic proteins can be found in Supplemental File 1.

*Supplemental Figure 3*

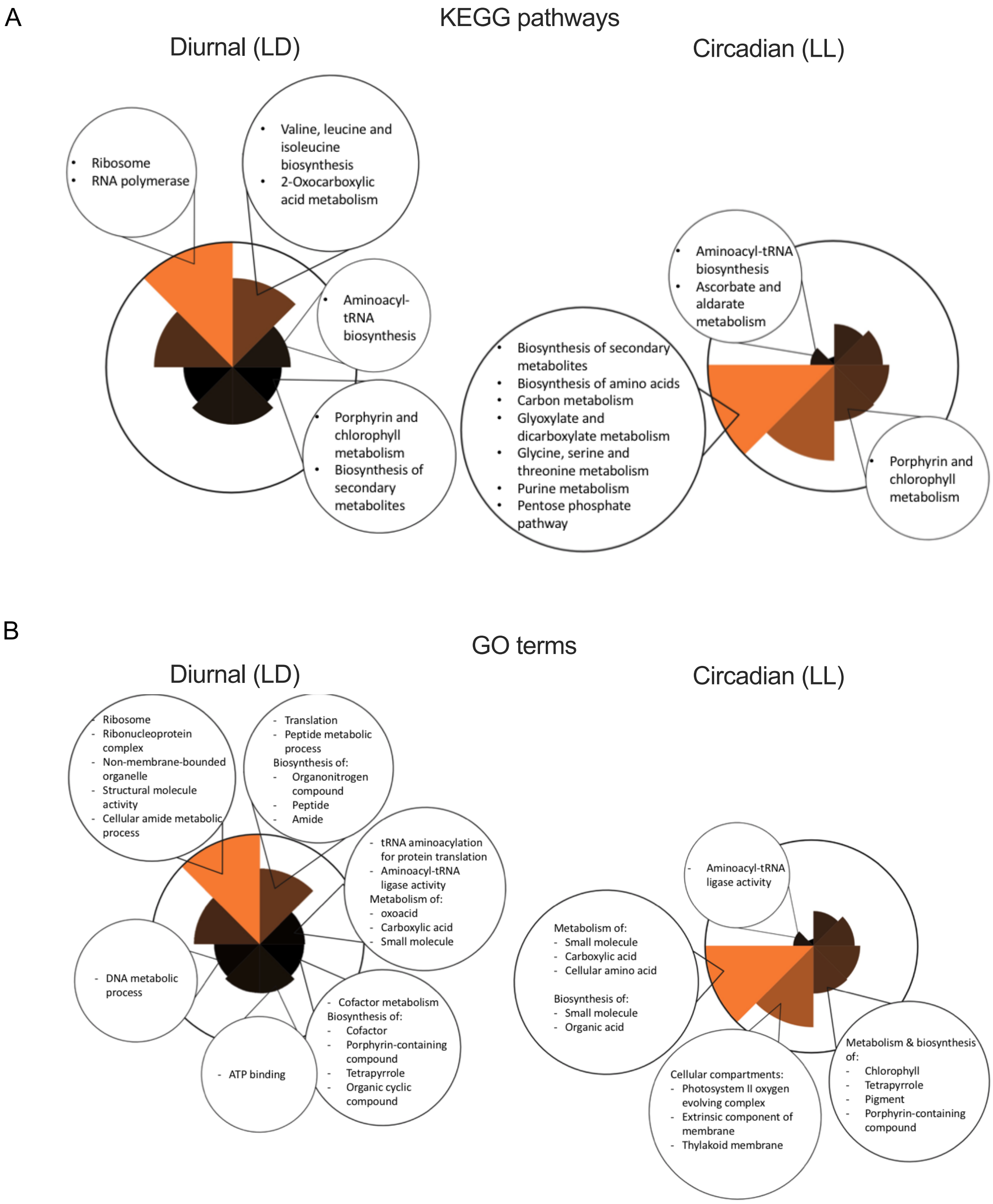


*Supplemental Figure 3: Enrichment of functional classifiers among protein peak phases*

A) KEGG processes that are significantly enriched amongst rhythmic proteins in 3-hour peak phase windows in LD (left) and LL (right). B) GO terms significantly enriched amongst rhythmic proteins in 3-hour peak phase windows in LD (left) and LL (right).

*Supplemental Figure 4*

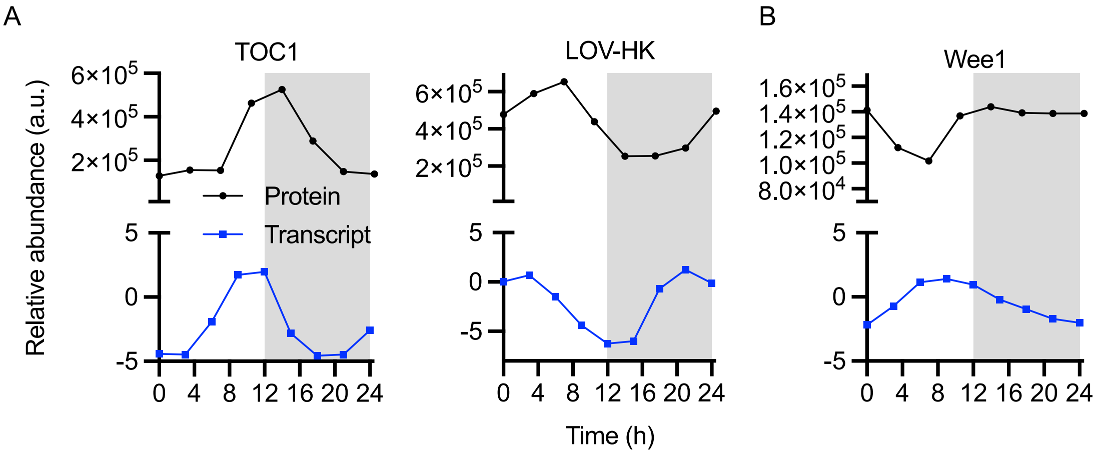


*Supplemental Figure 4: Relationships between transcript and protein phase under LD cycles*

A) The transcript and protein profiles of the clock protein TOC1 and the photoreceptor LOV-HK show clear phase harmonics, indicating a linear relationship. B) The transcript and protein profiles of the critical cell cycle kinase Wee1 are close to antiphasic, indicating a lack of phase harmonics.

*Supplemental Figure 5*

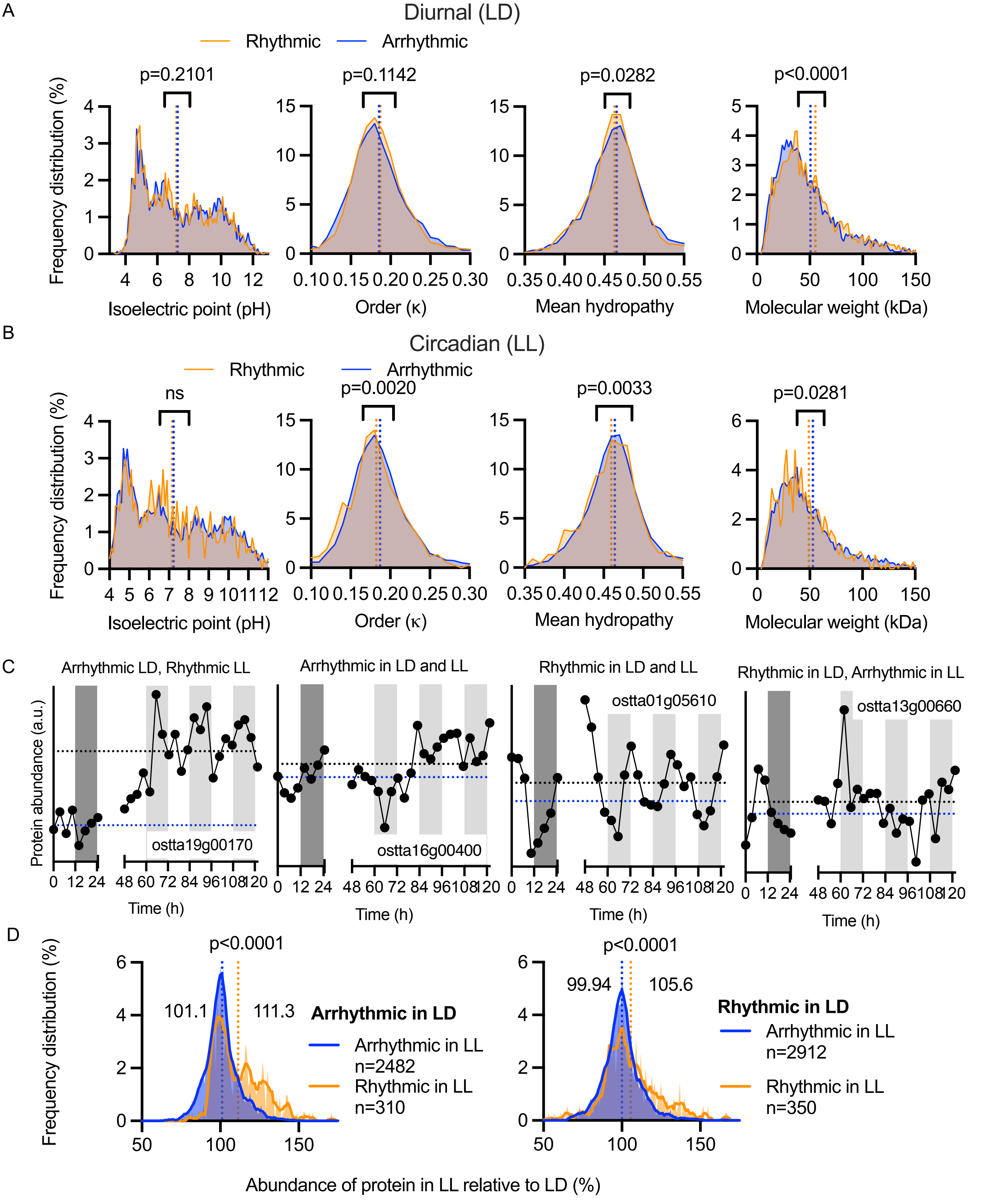


*Supplemental Figure 5: Biochemical properties of rhythmic versus arrhythmic proteins*

A-B) The biochemical properties of rhythmic (orange) versus arrhythmic (blue) proteins under entrained (A) or constant (B) conditions. Statistics reflect Mann-Whitney tests (ns = p>0.05). C) Example traces for each of the four combinations of rhythmic or arrhythmic in LD or LL, indicating the overall trend in main Fig. 4. Blue dotted lines are mean abundance under LD, black dotted lines mean abundance under LL. D) Frequency distribution of the mean abundance of rhythmic versus arrhythmic proteins in LL relative to their abundance in LD, split between proteins that are arrhythmic in LD (left) or those that are rhythmic in LD (right). Statistics reflect Mann-Whitney tests.

*Supplemental Figure 6*

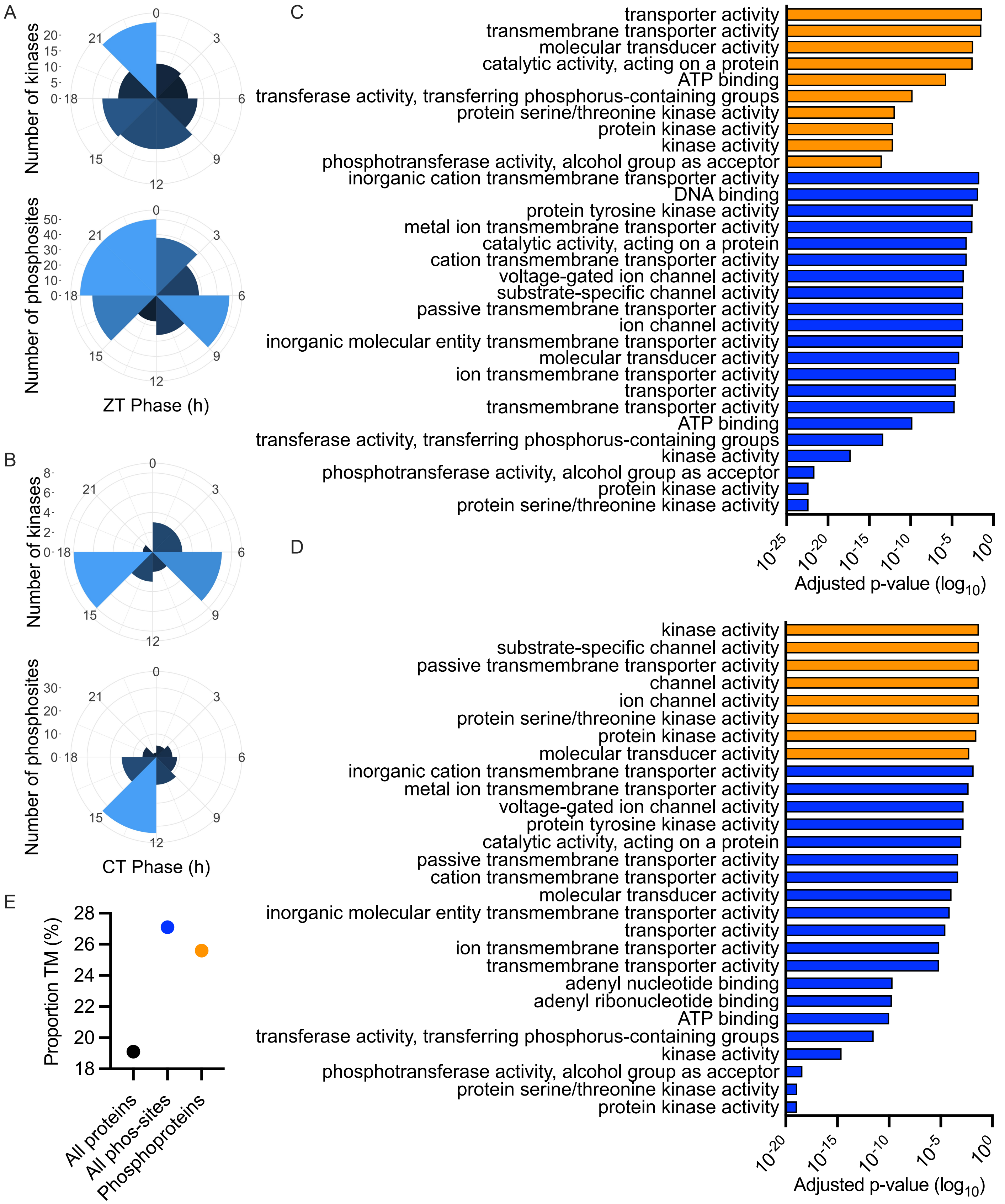


*Supplemental Figure 6: Rhythmic kinome and phosphoproteome regulate the transmembrane proteome*

A-B) Circular histograms showing the number of rhythmic kinases (top) and phosphosites (bottom) in LD (A) or LL (B) at each 3-hour peak phase interval. No phospho-enrichment was performed; these data represent phosphopeptides detected in the original TMT runs. C-D) Gene Ontology enrichment analysis for molecular function of all phosphorylated proteins (blue) or rhythmically phosphorylated proteins (orange) under LD (C) or LL (D). Under both conditions, membrane-associated functions (i.e. transporter activity) are overrepresented not only in phosphoproteins but also in rhythmically regulated phosphoproteins. E) The percentage of TM proteins among proteins carrying detected phosphorylation sites (blue) and among unique phosphoproteins (orange) is higher than among the overall detected proteome (black).

*Supplemental Figure 7:*

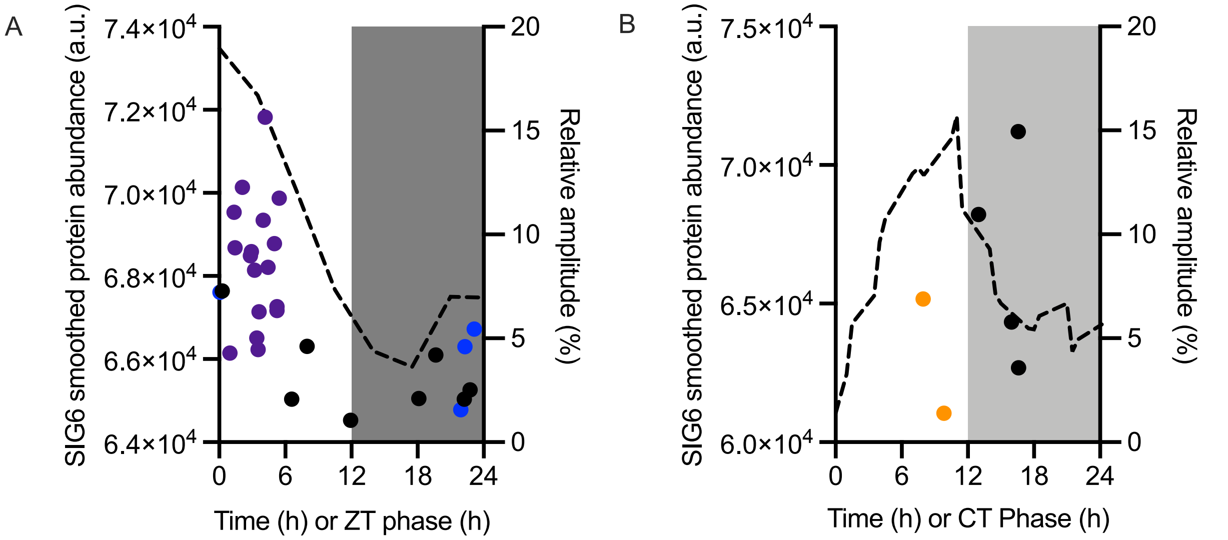


*Supplemental Figure 7: Phase relationship between the nuclear-encoded plastid transcription initiation factor SIG6 and the chloroplast encoded proteome*

Smoothed line of the mean abundance of SIG6 (left y-axes) under LD (A) or LL conditions (B) in relation to the amplitude (right y-axes) and peak phase of all rhythmic chloroplast-encoded proteins. Purple dots represent ribosomal proteins, blue RNA polymerases, and orange photosystem components.
